# An activator of a two-component system controls cell separation and intrinsic drug resistance in *Mycobacterium tuberculosis*

**DOI:** 10.1101/2025.07.16.665149

**Authors:** Liam D. McDonough, Shuqi Li, Vanisha Munsamy-Govender, Celena M. Gwin, Jeremy M. Rock, E. Hesper Rego

## Abstract

Unlike commonly studied rod-shaped bacteria, mycobacteria grow from their poles, requiring precise coordination between division and initiation of new pole growth. The mechanisms that mediate this transition are largely unknown, but likely represent a rich source of drug targets for the treatment of mycobacterial infections, including tuberculosis. Here, we identify TapA (MSMEG_3748/Rv1697) as a key regulator of this transition. TapA interacts with the sensor kinase MtrB at the septum to initiate a signaling cascade that ultimately results in the expression of the essential peptidoglycan hydrolases RipAB, amongst others, at the end of division. Loss of TapA disrupts division, dysregulates pole formation, and sensitizes *Mycobacterium tuberculosis* and other mycobacteria to several first and second-line TB antibiotics, establishing TapA as a potential therapeutic target, and defining a new link between cell cycle progression, envelope remodeling, and intrinsic antibiotic resistance in mycobacteria.

## INTRODUCTION

A major goal in the field of microbiology is to develop an understanding of the molecular mechanisms that allow bacteria to reproduce. These mechanisms provide a rich source of drug targets for the treatment of infectious diseases, and first-line treatments for several infections include inhibitors of the macromolecular complexes required for growth and division. While great progress has been made in understanding how model bacteria like *Escherichia coli*, *Bacillus subtilis*, and *Caulobacter crescentus* grow and divide, many bacterial pathogens deviate significantly from these model organisms. Importantly, the major human pathogen *Mycobacterium tuberculosis* and related Actinobacteria grow by building new envelope at their poles, rather than along their side walls. In polar-growing organisms, the site of division will become the site of new pole growth. Therefore, the timing between completion of cytokinesis and onset of new pole growth must be controlled by mechanisms that do not exist in lateral-growers, in which division and growth are spatially segregated. Here, we identify a mycobacterial protein, MSMEG_3748/Rv1697, that is necessary for the timing of cell separation in *Mycobacterium smegmatis* and *Mycobacterium tuberculosis*. Cells lacking MSMEG_3748/Rv1697 form chains of cells with unresolved septa. Importantly, *rv1697* is predicted to be essential in *M. tuberculosis* [1], and its depletion sensitizes the pathogen to a wide range of antibacterials, making this protein a promising drug target.

The molecular function of MSMEG_3748/Rv1697 has not been studied in mycobacteria, but the putative corynebacterium homolog, named SteA, is involved in cell separation in *Corynebacterium glutamicum*. SteA forms a membrane-bound complex with protein encoded in the same operon, SteB, which directly activates the peptidoglycan hydrolase RipA at the end of division [2, 3]. Here, we show that in mycobacteria, MSMEG_3748/Rv1697 operates differently than its putative homolog in corynebacteria. Instead of recruiting SteB (named MctB in mycobacteria), MSMEG_3748/Rv1697 interacts directly with the sensor kinase MtrB and promotes phosphorylation of the cognate response regulator MtrA. Phosphorylated MtrA then activates transcription of several genes important for intrinsic drug resistance, including the septal peptidoglycan hydrolases *ripA* and *ripB*, at the end of the cell cycle. Thus, we propose naming this protein <u>T</u>wo-component <u>A</u>ctivating <u>P</u>rotein A, TapA.

## RESULTS

### TapA, but not MctB, is required for cell separation in *M. tuberculosis and M. smegmatis*

To identify novel mycobacterial divisome proteins, we performed a co-immunoprecipitation of FLAG-tagged PonA1, the sole essential bifunctional PBP in *M. smegmatis* [4]. Amongst the hits were several expected proteins (Table S1), including PknB, a kinase that phosphorylates PonA1 [4, 5], as well as the sole serine/threonine phosphatase PstP [6]. Likewise, we detected enzymes involved in peptidoglycan synthesis or remodeling, including MurJ [7] and DacB [8]. We also detected several actinobacteria-specific proteins, including MSMEG_0317/PgfA, which we recently identified as a key protein involved in polar growth [9], and MSMEG_3748/TapA, a protein of unknown function in mycobacteria.

TapA has not been studied in mycobacteria, but it and the protein encoded in the same operon, MctB, are encoded in a region of the genome that shares high synteny with corynebacteria and other Actinobacteria. TapA and MctB have moderate amino acid identity with proteins encoded in this region in *C. glutamicum*, which are named SteA and SteB (41% and 29%, respectively) because *C. glutamicum* lacking these proteins exhibits increased <u>S</u>ensitivity <u>T</u>o <u>E</u>thambutol (Figure 1a) [2]. In *C. glutamicum*, SteA forms a complex with SteB, which directly activates the peptidoglycan hydrolase RipA to mediate the final stages of cell separation. Accordingly, deletion of either *steA* or *steB* results in *C. glutamicum* cell chaining [2, 3]. Thus, we asked if TapA and MctB are involved in cell separation of *M. tuberculosis*. As *tapA* (*rv1697*) is predicted to be essential in *M. tuberculosis* [1], we used CRISPRi to inhibit its expression and stained the resulting cells with a fluorescent D-amino acid to visualize cell morphology. After five days of depletion, we observed long chains of cells with unresolved septa, suggesting that the final stages of cell separation are impaired (Figure 1b). To quantify the morphological changes, we measured the area of individual cells or chains (Figure 1d) and their septum number (Figure 1e). We found that depleting *tapA* leads to cells with a large number of unresolved septa compared to wild-type. Importantly, the division defect could be complemented by expression of a recoded *tapA* that was insensitive to CRISPRi depletion (Figure 1b,d-e). We also observed occasional bulging or lysed cells trapped within a chain, suggestive of a cell wall defect. Thus, as with SteA in *C. glutamicum*, TapA is required for cell separation in *M. tuberculosis*. Next, we asked if *tapA*’s operon partner, *mctB*, which plays a role in copper resistance [10], is also involved in *M. tuberculosis* cell separation. We designed two separate single-guide RNAs (sgRNAs) targeting the *mctB* open reading frame; surprisingly, expression of either *mctB*-targeting sgRNAs produced cells with no change in cell size or shape as compared to a non-targeting sgRNA control (Figure 1b, d-e). Together, these results suggest that *tapA*, but not its operon partner *mctB/steB*, is required for proper cell separation in *M. tuberculosis*.

**Figure 1:**
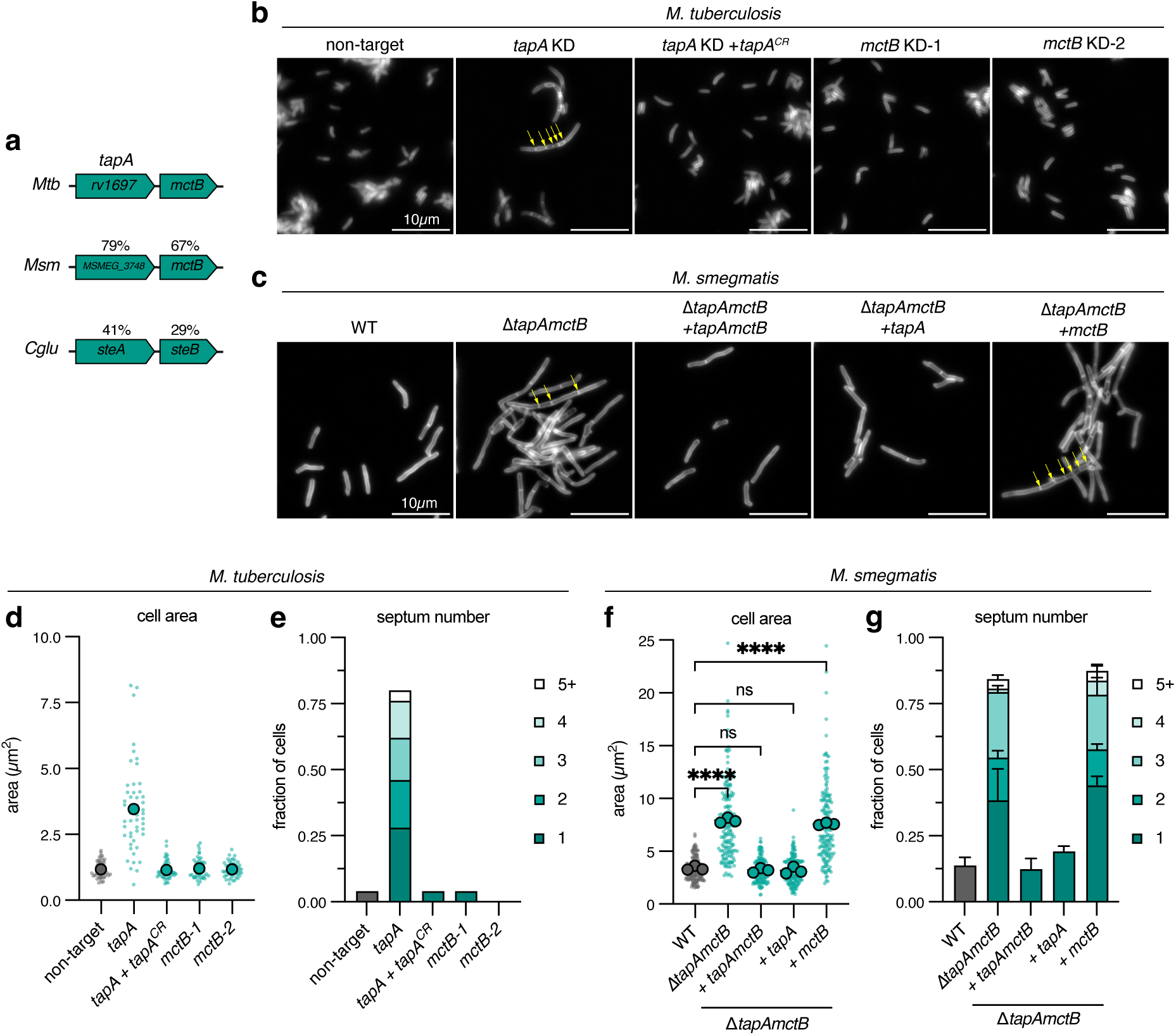
TapA, but not MctB, is required for cell separation in *M. tuberculosis* and *M. smegmatis*. **a**, Operon configuration of *M. tuberculosis* genes *rv1697*/*tapA* and *mctB* and amino acid identity of *M. smegmatis* and *C. glutamicum* sequence homologs. **b**, FDAA-stained *M. tuberculosis* CRISPRi strains after 5 days CRISPRi depletion. Yellow arrows indicate chains of unresolved septa. KD, knockdown; CR, CRISPRi-resistant. Two independent sgRNAs targeting *mctB* were tested (KD-1 and KD-2). Scalebar = 10µm. **c**, FDAA-stained exponential-phase *M. smegmatis* mutant and complement strains. Yellow arrows indicate chains of unresolved septa. Scalebar = 10µm. **d-e**, *M. tuberculosis* cell area and septum number quantification. Representative data from one of two independent experiments; *n* = 50 cells. Large dots indicate means, and small dots indicate individual cells. Septum number expressed as fraction of total cells. **f-g**, *M. smegmatis* cell area and septum number quantification; *N =* 3 independent experiments; *n* (cells) = 50, 68, 49 (WT); *n* = 56, 65, 52 (Δ*tapAmctB*); *n* = 48, 60, 60 (Δ*tapAmctB + tapAmctB*); *n* = 57, 61, 63 (Δ*tapAmctB + tapA*); *n* = 68, 52, 55 (Δ*tapAmctB + mctB*). Large dots indicate area means, and small dots indicate individual cells. Statistical significance was determined by ordinary one-way ANOVA followed by Dunnett’s multiple comparisons test on means of cell area data. ns = not significant (p > 0.05), * p < 0.05, ** p < 0.01, *** p < 0.001, **** p < 0.0001. Septum number expressed as fraction of total cells.

To understand the role of TapA and MctB/SteB in cell separation of other mycobacteria, we attempted to produce a *tapAmctB* deletion mutant by recombineering in *M. smegmatis* [11]. We obtained mutants on agar plates but were unable to propagate these as a turbid culture in our typical growth medium (Figure S1a). Some cell-wall-defective mutants can be grown in osmoprotective medium that contains high concentrations of sucrose, magnesium chloride, and maleate (SMM) [12]. Indeed, the addition of SMM to the medium allowed for the stable propagation of Δ*tapAmctB* (Figure S1a), but cells remained elongated and multiseptated (Figure 1c). This allowed us to assess the individual role of *tapA* or *mctB* in cell separation by complementation of the double mutant with either single gene or the full operon using an integrating vector. Consistent with our results in *M. tuberculosis*, complementation with the full operon or *tapA* alone produced cells with wild-type size and septum number, while complementation with *mctB* alone phenocopied the parental deletion mutant (Figure 1f,g). Thus, as in *M. tuberculosis*, *tapA*, but not its operon partner *mctB*, is required for cell separation in *M. smegmatis*.

In *C. glutamicum*, SteB contributes to robust recruitment of SteA to the septum [2]. To test if MctB is involved in TapA septal localization, we expressed GFP-tagged alleles of either *M. smegmatis* or *M. tuberculosis tapA* in Δ*tapAmctB M. smegmatis*. Expression of either allele from an integrating vector restored cell separation, and cells exhibited septal GFP localization that was unaltered by the presence or absence of the cognate *mctB* gene (*MSMEG_3747* or *rv1698*, respectively) (Figure S1b,c). In contrast, *C. glutamicum* SteA localized more strongly to the *M. smegmati*s septum in the presence of SteB, consistent with the reported results in *C. glutamicum* [2], but expression of neither *steA* nor *steAB* restored cell separation in Δ*tapAmctB M. smegmatis* (Figure S1d). Together, these data show that TapA, unlike SteA in *C. glutamicum*, is not recruited to the septum by MctB but is still required for cell separation in both *M. tuberculosis* and *M. smegmatis*.

### TapA genetically interacts with the MtrAB two-component system, and operates in the same pathway

Our data indicate that TapA’s essential function is independent of MctB/SteB in mycobacteria, consistent with recent data suggesting a SteB-independent function of SteA in *C. glutamicum* [13]. To uncover this function, we employed unbiased screens to identify genetic interactors in both *M. tuberculosis* and *M. smegmatis*. First, to discover *tapA* genetic interactions in *M. tuberculosis*, we analyzed data from a genome-wide chemical-genetic CRISPRi screen we previously performed, which profiled the fitness of depletion mutants exposed to various anti-TB compounds [14]. Similar signatures in chemical-genetic screens can uncover genes operating in the same pathway [14–16]. Analysis of the chemical-genetic data revealed that gene knockdown of *tapA* closely resembled knockdown of the two-component system *mtrAB*, along with MtrA-regulated genes *ripA* and *pirG* (Figure 2a). To confirm our pooled approach, we constructed strains expressing individual sgRNAs targeting each gene (Figure 2b). Pre-depletion of *tapA*, *mtrA*, or *mtrB* led to strong sensitization to rifampin, vancomycin, and bedaquiline, as compared to a control non-targeting sgRNA, recapitulating the genome-wide screen. We observed similar multi-drug sensitization for *tapA* mutants in *M. smegmatis* (Table S3). In contrast to deletion of *steA* in *C. glutamicum* [2], pre-depletion or deletion of these genes did not greatly alter sensitivity to ethambutol in either *M. tuberculosis* or *M. smegmatis* (Figure 2b and Table S3). Rifampin, bedaquiline, and vancomycin all employ different mechanisms of action, but are large hydrophobic drugs that do not easily cross the mycobacterial cell envelope, suggesting that depletion of these genes disrupts the cell envelope in similar ways. These similar chemical-genetic signatures suggest a functional relationship between TapA and the MtrAB two-component system.

**Figure 2:**
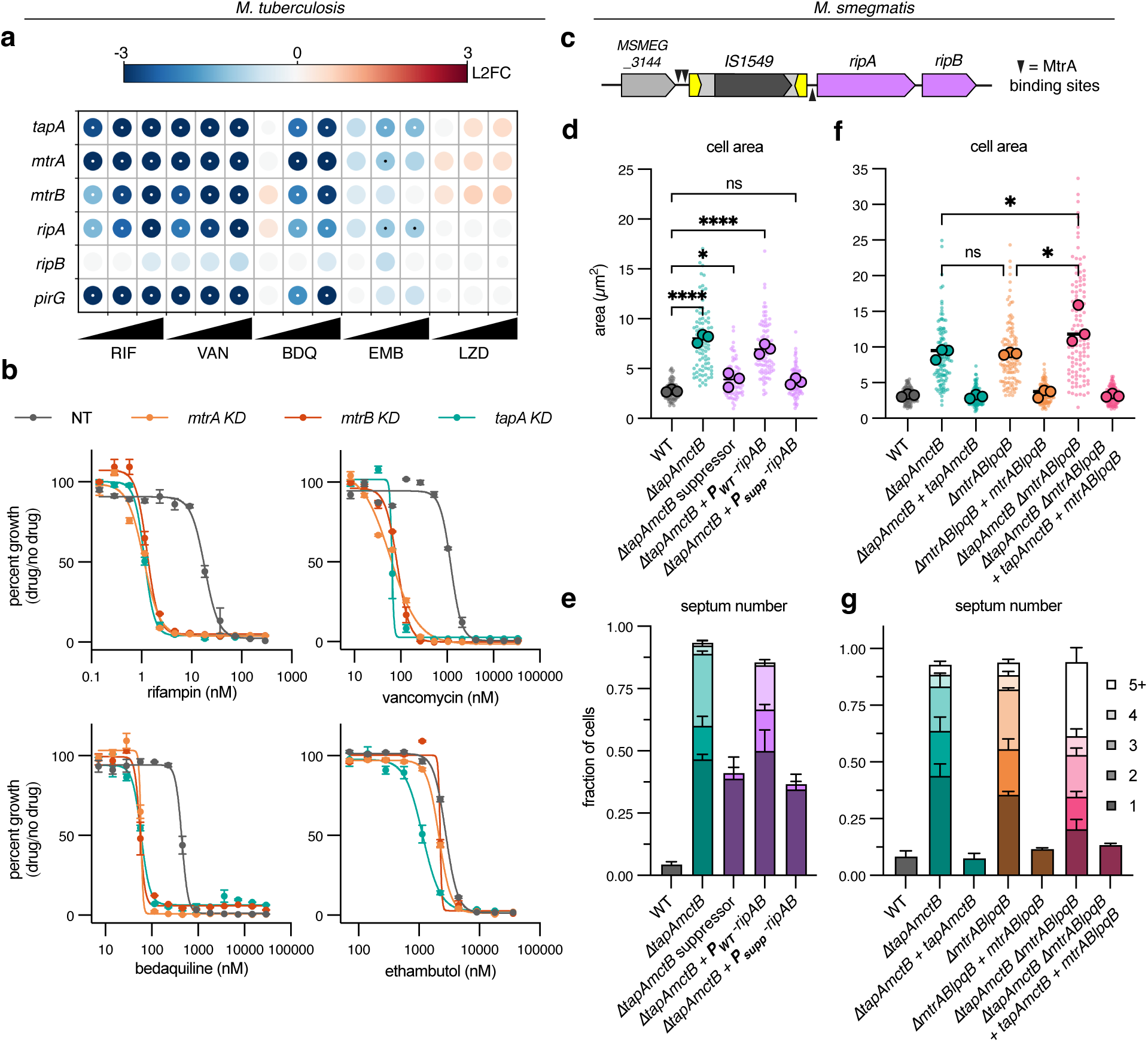
Unbiased screens reveal a functional relationship between TapA and the MtrAB two-component system. **a**, Feature-expression heatmap of *tapA*, *mtrAB*, and MtrA-regulated genes *ripAB* and *pirG* from a 5 day CRISPRi library pre-depletion screen. The color of each circle represents the gene-level log2 fold change (L2FC); a white or black dot represents a false discovery rate (FDR) <0.01 and a |L2FC|>1. VAN = vancomycin; RIF = rifampicin; BDQ = bedaquiline; EMB = ethambutol. Each antibiotic was tested in triplicate at three sub-minimum inhibitory concentrations (sub-MIC90) listed in Table S2. **b**, Dose-response curves (mean ± s.e.m., *n* = 3 biological replicates) for the indicated *M. tuberculosis* strains expressing individual sgRNAs. **c**, Genomic organization of a suppressor mutant in *M. smegmatis ΔtapAmctB* containing an insertion sequence (*IS1549*) in the promoter of the cell wall hydrolase operon *ripAB*. **d-e**, Cell area and septum number quantification of *M. smegmatis* strains from screen and strains expressing single copies of cloned *ripAB* promoters from WT or suppressor mutant driving *ripAB* expressing; *N =* 3 independent experiments; *n =* 30 cells per replicate. **f-g**, Cell area and septum number of *M. smegmatis* single and double operon mutants. *N =* 3 independent experiments; *n* (cells) = 31, 43, 50 (WT); *n* = 39, 31, 53 (Δ*tapAmctB*); *n* = 34, 47, 46 (Δ*tapAmctB + tapAmctB*); *n* = 32, 37, 52, (Δ*mtrABlpqB*); *n =* 38, 40, 51 (Δ*mtrABlpqB + mtrABlpqB*), *n* = 41, 38, 40, (Δ*tapAmctB ΔmtrABlpqB*); *n* = 34, 45, 50 (Δ*tapAmctB ΔmtrABlpqB* + *tapAmctB* + *mtrABlpqB*). Large dots indicate area means, and small dots indicate individual cells. Statistical significance was determined by ordinary one-way ANOVA followed by Tukey’s multiple comparisons test on means of cell area data. Selected comparisons between single and double operon mutants are shown for clarity. ns = not significant (p > 0.05), * p < 0.05, ** p < 0.01, *** p < 0.001, **** p < 0.0001. Septum number expressed as fraction of total cells.

Second, to identify genetic interactors of *M. smegmatis tapA*, we designed a suppressor mutant screen. In our initial attempts to culture *M. smegmatis* Δ*tapAmctB* cells, we found that supplementation of media with 100mM sucrose alone was enough to sustain growth without producing a fully turbid culture (Figure S2a). We reasoned that this intermediate growth condition would allow for selection of suppressor mutants that grow turbidly. To this end, we passaged eight independent populations of Δ*tapAmctB* cells in sucrose-supplemented media until turbid mutants arose after about 20 days (Figure S2b). To enrich for turbidly growing mutants, we next passaged the sucrose-evolved populations in no-sucrose media, a non-permissive condition for the original parent strain (Figure S2c), and plated for single colonies. After confirming that single-colony isolates grew turbidly, we stained exponential phase cultures with a fluorescent D-amino acid to check if suppression of the macroscopic culture growth defect also suppressed the cell chaining. While most isolates remained as chains (Figure S2d), one serendipitous mutant restored normal septum number (Figure S2e). Whole genome sequencing of this mutant revealed that an endogenous insertion sequence, *IS1549* [17], had inserted in the promoter region of *ripAB*, an operon encoding two peptidoglycan hydrolases that are together essential for cell separation [18, 19] (Figure 2c, Table S4). The transposon inserted such that it displaced reported binding sites for the response regulator MtrA, which controls *ripAB* expression [20]. To confirm that this insertion sequence was responsible for suppressing the chaining defect of *ΔtapAmctB* cells, we cloned an integrating vector expressing *ripAB* from its endogenous promoter, with or without the insertion sequence. Only the *IS1549-*containing promoter could restore normal septum number and area in the parental Δ*tapAmctB* strain (Figure 2d-e). Together with the *M. tuberculosis* chemical-genetic interactions, these data pointed to a functional relationship between TapA and the expression of the peptidoglycan hydrolases *ripAB*, mediated by the MtrAB two-component system.

The histidine kinase MtrB localizes to the septum, is activated by unknown signals, and phosphorylates its cognate response regulator MtrA. MtrA∼P then promotes expression of *ripAB* and several other genes involved in cell division, DNA replication, and antibiotic sensitivity [20]. Consequently, deletion or depletion *mtrA*, *mtrB*, or *ripA* produces chained cells [18–20]. However, while most histidine kinases sense environmental signals to regulate gene expression [21], it is not known how MtrB is activated to ensure proper expression of its regulon, including *ripAB*, at the final stages of cell division. To test if TapA is involved in the MtrAB signaling cascade, we constructed an *mtrABlpqB* deletion mutant and a double operon mutant lacking *tapAmctB* and *mtrABlpqB* in *M. smegmatis*. Deletion of either operon resulted in comparable increases in cell size and septum number, which could be complemented by expression of the corresponding genes on an integrating vector (Figure 2f-g). Cells lacking all five genes had a slightly higher average size and septum number (Figure 2f-g); nevertheless, the combined effect was much less dramatic than deleting either operon individually, as compared to wild-type cells. Likewise, the double operon mutant had similar antibiotic sensitivity to mutants lacking each operon individually (Table S3). Thus, while TapA and MtrAB may have small independent roles in cell separation, these results, together with results from the unbiased screens described above, led us to hypothesize that TapA is a positive regulator of the MtrAB two-component system, which activates the expression of *ripAB*.

### TapA and MtrB physically interact and co-localize at the septum at the end of division

To test our hypothesis, we first asked if TapA was physically interacting with a member of the MtrAB-LpqB “three-component system” [22] or other potential unknown proteins involved in cell separation. We performed a co-immunoprecipitation in *M. smegmatis* expressing msfGFP-TapA from its native promoter as the sole copy in the cell. This strain grows with normal morphology, indicating the msfGFP tag does not disrupt TapA’s function (Figure S1b). To help identify TapA-specific interactions, we simultaneously pulled down other msfGFP-tagged proteins involved in cell growth and division, including FtsZ, Wag31, and PgfA, among others, and searched for peptides that were more abundant in the TapA precipitates (Table S5). We used a GFP nanobody to precipitate msfGFP from cell lysates and identified co-precipitating peptides by mass spectrometry. Amongst the peptides most enriched in the TapA pulldown, but not others, were MtrB and its associated lipoprotein LpqB, as well as several other histidine kinases. MtrB was also highly enriched in the Wag31 precipitates, consistent with a previously published study showing that MtrB interacts with both Wag31 and FtsI [23]. To validate the TapA-MtrB interaction in an orthogonal system, we used a bacterial adenylate cyclase two-hybrid assay (BACTH) (Figure 3a) [24]. In an *E. coli* strain lacking its endogenous adenylate cyclase, proteins of interest were fused to two fragments, T25 and T18, of the *Bordetella pertussis* adenylate cyclase. Interaction between proteins of interest will reconstitute the adenylate cyclase, producing cAMP that drives expression of a reporter β-galactosidase, which can be assessed qualitatively on X-gal plates or quantitatively by an ortho-nitrophenyl-β-galactoside (ONPG)-based assay. As a positive control, we co-expressed T25-MtrB and T18-MtrA and observed blue color on X-gal plates and a positive signal in the ONPG assay. Consistent with the co-IP results, production of T25-MtrB and T18-TapA—or, reciprocally, T18-MtrB and T25-TapA—showed interaction by both assays (Figure 3a), while TapA did not show an interaction with the response regulator MtrA (Figure S3a).

**Figure 3:**
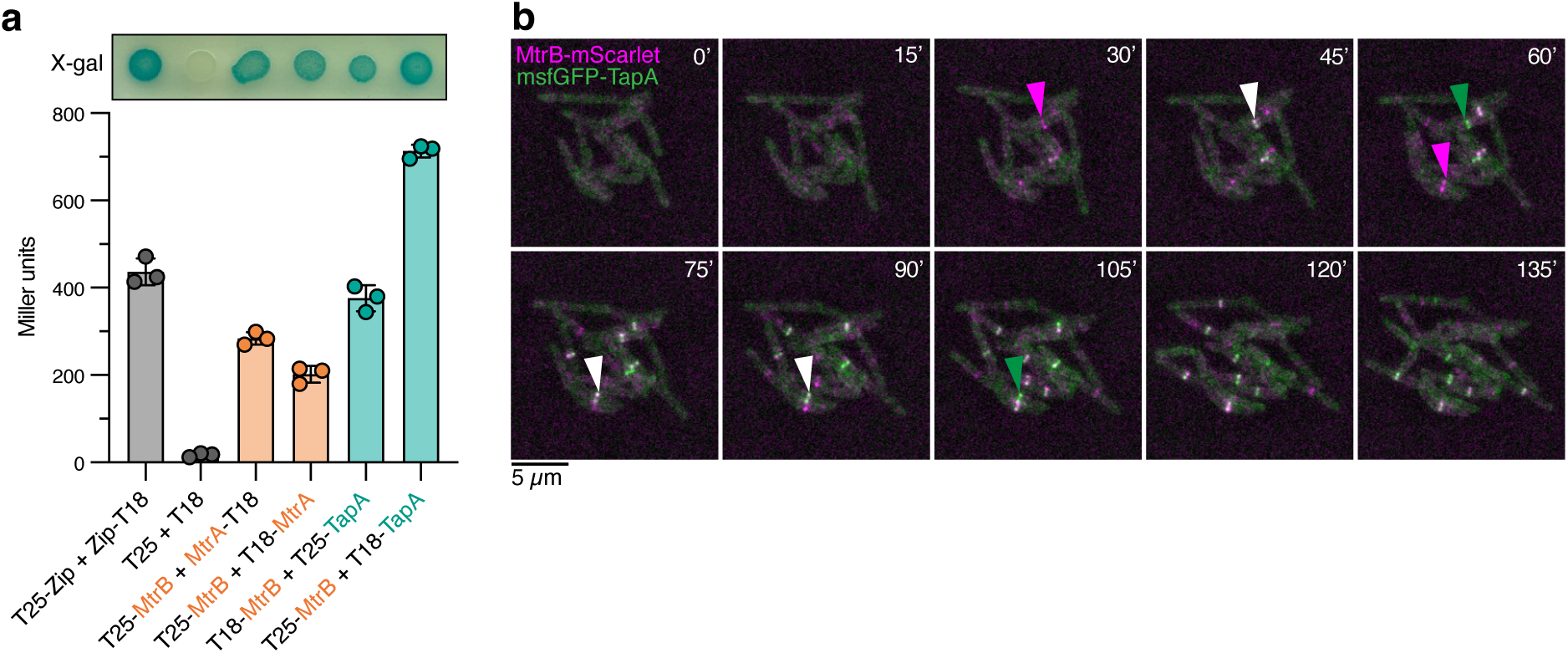
TapA and MtrB interact and colocalize at the end of the cell cycle. **a**, Bacterial two-hybrid assays after 43 hours post-induction with IPTG on X-gal plates (top) or in buffer containing ONPG (bottom). **b**, Montage of timelapse microscopy of *M. smegmatis* expressing MtrB-mScarlet (magenta arrowheads) and msfGFP-TapA (green arrowheads) as the sole gene copies. White arrowheads point to colocalized proteins.

An Alphafold3 prediction of a TapA homodimer produced a moderate-confidence structure (Figure S3b). The placement of the TapA globular domains relative to the transmembrane helices suggested that TapA would likely interact with the transmembrane and HAMP domains of MtrB. To test this, we constructed an MtrB truncation mutant (MtrB_1-285_) lacking the kinase domain and found that it also interacts with TapA in the bacterial two-hybrid assay (Figure S3a). Together, these results confirmed that TapA physically interacts with MtrB and showed that the kinase domain of MtrB is not required for this interaction.

Since TapA and MtrB physically interact, we reasoned that they would co-localize at the septum. To visualize their localization, we constructed an *M. smegmatis* strain expressing msfGFP-TapA and MtrB-mScarlet as the sole copies of each gene in the cell, each driven from their respective native promoter sequences. To track the localization of TapA and MtrB over the course of the cell cycle, we performed time-lapse phase contrast (Figure S3c) and fluorescence microscopy (Figure 3b) in a microfluidic chamber with constant perfusion of media. This strain has wild-type morphology, indicating each tagged allele is functional, and we observed bands of both msfGFP-TapA and MtrB-mScarlet localization at the septum (Figure 3b). MtrB-mScarlet arrives at the septum first, then msfGFP-TapA colocalizes for several frames, followed by dissipation of the MtrB-mScarlet signal. The msfGFP-TapA signal persists until the cell divides. Together with the pulldown and bacterial two-hybrid data, these results show that MtrB and TapA physically interact at the septum near the end of the cell cycle to mediate cell separation.

### TapA is required for MtrA-dependent gene expression

We hypothesized that TapA could be involved in recruiting or stabilizing MtrB to the septum and/or in activating the MtrAB phospho-relay. To test if MtrB and TapA depend on one another for proper localization at the septum, we compared the localization of each protein in the absence of the other. MtrB-mScarlet localizes to the septum in a Δ*tapAmctB* strain, and msfGFP-TapA localizes to the septum in a Δ*mtrABlpqB* strain (Figure S3d), indicating that neither protein is required for recruitment of the other to the septum. Thus, other unknown recruitment mechanisms must be responsible for the ordered arrival of these proteins at the septum.

To test if TapA is required for expression of the MtrAB regulon, we measured the relative abundance of MtrA-regulated transcripts by RT-qPCR. In the *M. tuberculosis* CRISPRi strains, depletion of *mtrA*, *mtrB*, or *tapA* resulted in reduced *ripA*, *ripB*, and *rpfC* expression relative to the housekeeping gene *sigA*, as compared to the non-targeting control strain (Figure 4a). Knockdown of *tapA* did not alter expression of *mtrA* or *mtrB* (Figure 4a), and expression of *rpoB* was also unaltered by knockdown of these genes (Figure S4). Correspondingly, expression of *M. smegmatis* genes *ripA* and *pirG* was reduced in cells lacking *tapA* (Figure 4b-c). The reduction in *ripA* and *pirG* transcripts was relatively equal in Δ*tapAmctB* and *ΔmtrABlpqB* cells, or cells expressing the phospho-ablative mutants *mtrA-D56A* or *mtrB-H279Y*. These data are consistent with TapA being a positive regulator of the MtrAB two-component system.

**Figure 4:**
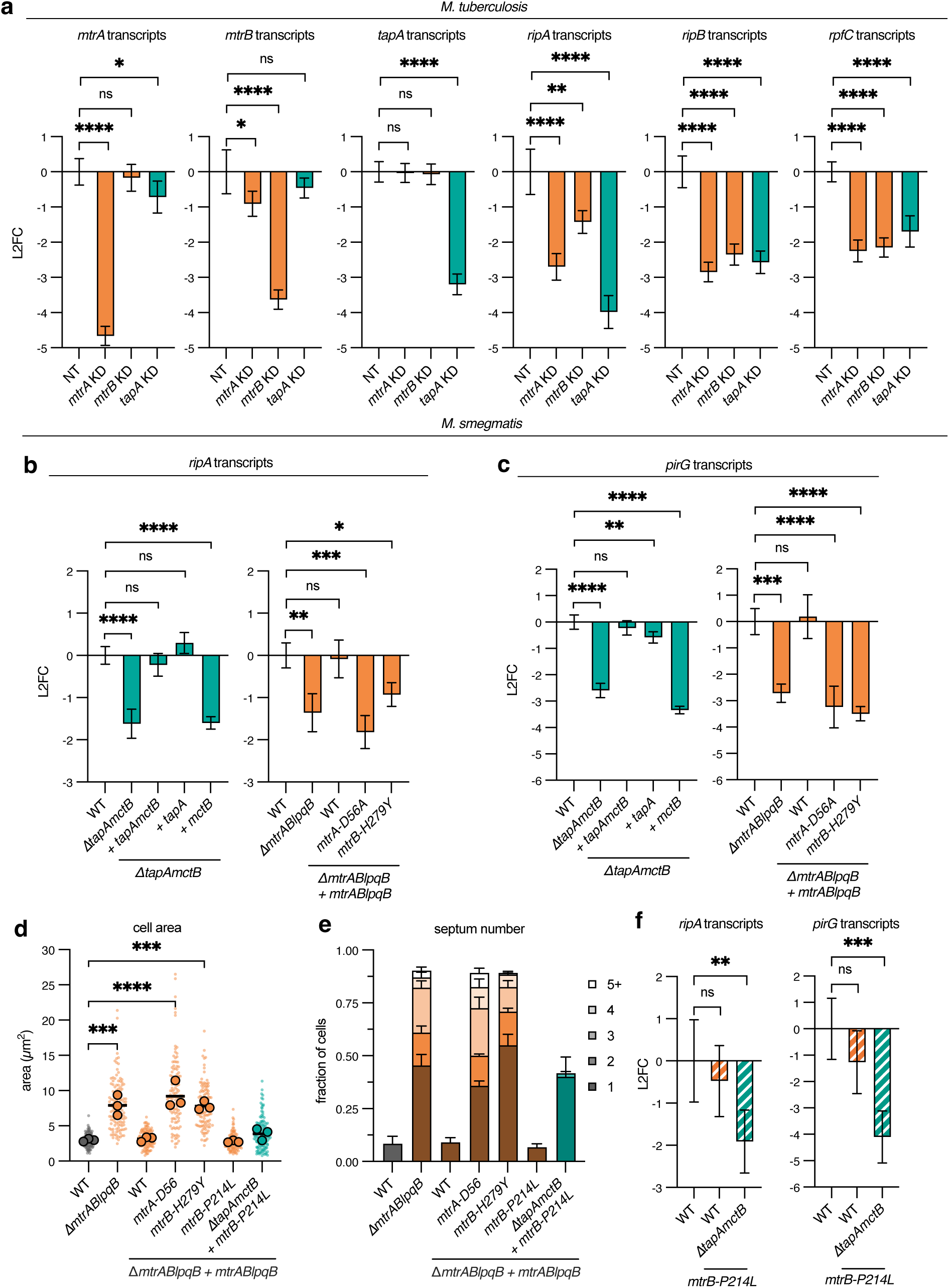
TapA is required for MtrA-dependent gene expression. **a**, *M. tuberculosis* transcripts in CRISPRi knockdown (KD) strains. KD target genes are *mtrA*, *mtrB*, *tapA*, and a non-targeting (NT) control. Transcript abundances for indicated genes are normalized to that of *sigA*, and expressed as L2FC of NT control. *n =* 3 biological replicates. **b-c**, *M. smegmatis* transcripts in the indicated mutant and corresponding complementation strains. Transcript abundances for *ripA* (**b**) and *pirG* (**c**) are normalized to that of *sigA*, and expressed as L2FC of WT control. *n =* 3 biological replicates. Statistical significance for **a-c** was determined by ordinary one-way ANOVA followed by Dunnett’s multiple omparisons test on underlying ΔCt values. ns = not significant (p > 0.05), * p < 0.05, ** p < 0.01, *** p < 0.001, **** p < 0.0001. **d-e**, Cell area and septum number quantification of *mtrABlpqB* complementation strains; *N =* 3 independent experiments; *n* (cells) = 40, 40, 40 (WT); *n* = 40, 41, 42 Δ*mtrABlpqB*); *n* = 40, 40, 41 (Δ*mtrABlpqB + mtrABlqpB* WT); *n* = 40, 40, 40 (Δ*mtrABlpqB + [mtrA-D56A]-mtrBlqpB*); *n* = 40, 40, 40 (Δ*mtrABlpqB + mtrA-[mtrB-H279Y]-lqpB*); *n* = 40, 40, 40 (Δ*mtrABlpqB + mtrA-[mtrB-P214L]-lqpB*); *n* = 43, 40, 43 (Δ*tapAmctB* Δ*mtrABlpqB + mtrA-[mtrB-P214L]-lqpB*). Large dots indicate area means, and small dots indicate individual cells. Statistical significance was determined by ordinary one-way ANOVA followed by Dunnett’s multiple comparisons test on means of cell area data. * p < 0.05, ** p < 0.01, *** p < 0.001, **** p < 0.0001. Significance for p > 0.05 is not shown for clarity. Septum number expressed as fraction of total cells. **f**, *M. smegmatis* transcripts in strains expressing *mtrB-P214L*. Statistical significance determined as in **a-c**.

We reasoned that if TapA’s essential role is mediated via MtrAB, then a gain-of-function MtrB mutation would bypass the loss of *tapA*. Several mutagenesis and directed evolution studies in other bacteria have uncovered loss– and gain-of-function mutations for histidine kinases [25–27]. In particular, mutations in the HAMP signal transduction domain can lead to increased phosphorylation of the kinase and subsequent target gene expression. Mutation of a highly conserved proline to leucine in the *hitS* histidine kinase in *Bacillus anthracis* results in constitutive kinase activity [27], and this proline is also conserved in *mtrB* homologs across mycobacterial species, as well as other proteins containing HAMP-domain (Figure S5). We constructed a strain expressing *mtrB-P214L* as its sole copy, in either wild-type or Δ*tapAmctB M. smegmatis*. Cells expressing this allele in an otherwise wild-type background were morphologically indistinguishable from wild-type cells for both cell area and septum number (Figure 4d-e). Consistent with our hypothesis, expression of *mtrB-P214L* partially rescued the loss of *tapA*, resulting in cells with near wild-type area and septum number (Figure 4d-e). However, 40% of Δ*tapAmctB mtrB-P214L* cells had one septum, as compared to 8% of wild-type cells, indicating only partial rescue of cell separation (Figure 4e). Expression of *mtrB-P214L* also did not fully restore *ripA* or *pirG* gene expression in Δ*tapA* cells (Figure 4f-g). Nevertheless, these data are consistent with TapA acting as an activator of MtrAB, leading to increased expression of the essential peptidoglycan hydrolases *ripAB*.

### TapA is required for MtrA phosphorylation

A key step in a two-component signal transduction pathway is the phosphorylation of the response regulator by the histidine kinase. Specifically, phosphorylation of MtrA by MtrB alters MtrA’s DNA-binding affinity, most often resulting in activation of target gene expression [20]. Since TapA is required for the expression of MtrA-regulated genes, we hypothesized that MtrA phosphorylation is dependent on TapA. To test this, we constructed a Myc-tagged allele of MtrA, again as the sole copy in the cell, and measured

MtrA phosphorylation in various strain backgrounds by Phos-tag^TM^ SDS-PAGE followed by quantitative Western blot (Figure 5d-e). Phos-tag gels resolve phosphorylated and dephosphorylated forms of proteins, allowing for quantification of phosphorylation in various strain backgrounds [28]. The Myc-MtrA allele produced wild-type cell morphology, indicating that it is functional (Figure S6). We confirmed that the upper band corresponds to phosphorylated MtrA, as it was undetectable in cell lysates expressing the phospho-ablative alleles *mtrA-D56A* and *mtrB-H279Y*. Across three independent experiments, approximately 10% of MtrA was phosphorylated in wild-type cells, consistent with MtrA’s role in cell division, as only a fraction of cells are actively separating at a particular time in an asynchronous culture. As in our RT-qPCR experiments, MtrA phosphorylation was reduced only in cells lacking *tapA*, but not *mctB*. Additionally, the *mtrB-P214L* gain-of-function mutation partially restored MtrA phosphorylation in the absence of *tapA* (from ∼1% to 5%). Overall, these data support the model that TapA activates phosphotransfer from MtrB to MtrA, potentially by inducing a conformational change in MtrB to its active kinase state. Furthermore, these data suggest that TapA is required for the proper timing of MtrA phosphorylation and subsequent cell separation.

**Figure 5:**
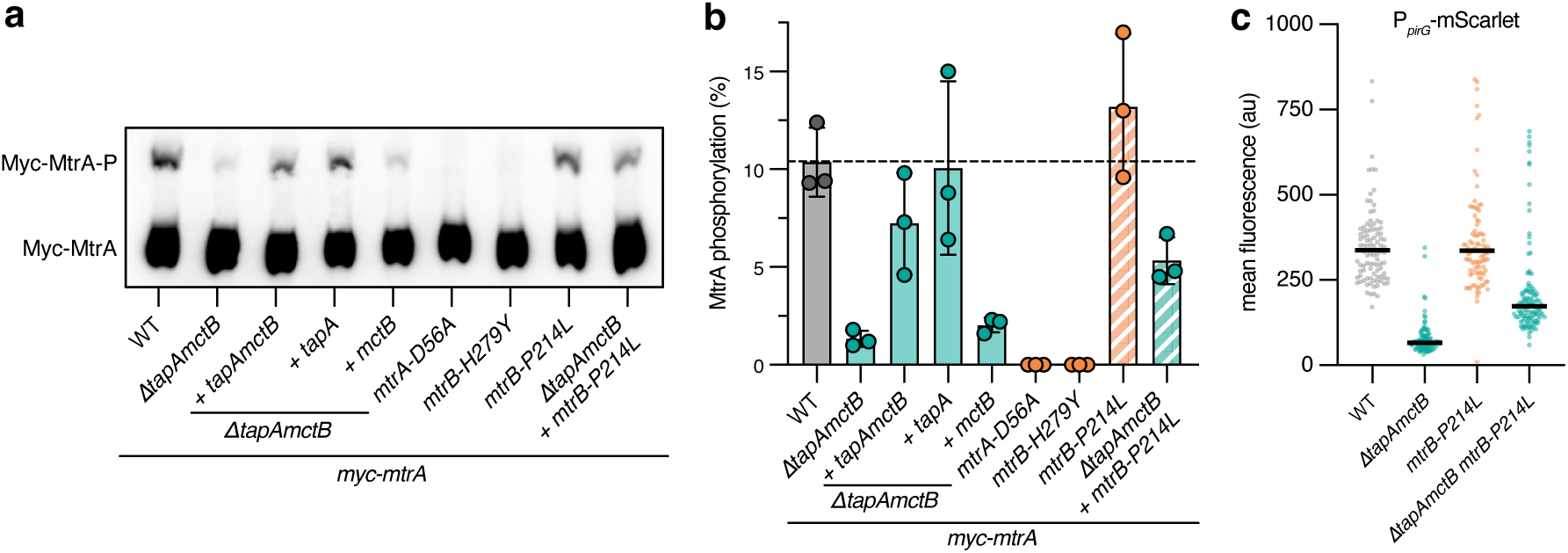
TapA is required for MtrA phosphorylation and full promoter activity of *pirG*. **a**, Representative Western blot following Phos-tag SDS-PAGE of cell lysates expressing *myc-mtrA* in the indicated strain backgrounds. Upper bands correspond to Myc-MtrA-P. **b**, Quantification of blots represented in **a**. MtrA phosphorylation is calculated as the fraction of total Myc-MtrA protein. *N* = 3 independent experiments. **c**, P*_pirG_*-mScarlet signal of individual cells from exponential phase cultures. WT, Δ*tapAmctB*, *mtrB-P214L*, or Δ*tapAmctB* + *mtrB-P214L* strains were grown in HdB-SMM. Mean fluorescence of individual cells and median of the population (black bars) are shown. Representative results from two independent experiments. *n* (cells) = 99 (WT); *n* = 97 (Δ*tapAmctB*); *n =* 100 (*mtrB-P214L*); *n* = 116 (Δ*tapAmctB + mtrB-P214L*).

### TapA mediates cell-cycle-dependent expression of MtrA-regulated genes

Together, these data lead to a model whereby TapA, through a direct protein-protein interaction with MtrB, activates the signal transduction cascade that leads to the expression of *ripAB* and other genes in the MtrA regulon at the end of cell division. This model predicts that MtrA-mediated gene expression occurs dynamically, peaking near the end of the cell cycle, in a TapA-dependent manner. To test this, we constructed transcriptional reporters for MtrA-regulated promoters fused to the fluorescent protein mScarlet-I to visualize single-cell gene expression dynamics. We first attempted to make a reporter driven by the promoter of *ripAB*, but were unable to observe fluorescence above background, consistent with these peptidoglycan hydrolases being expressed at low levels. Other genes in the MtrAB regulon are expressed at higher levels, including *pirG*, a gene involved in intrinsic drug resistance and cell envelope integrity across multiple mycobacterial species (Figure 2a) [14, 29, 30]. We cloned the native promoter sequence of *M. smegmatis pirG* to drive expression of mScarlet on an integrating vector, then imaged individual cells from exponential-phase cultures (Figure 5c). Deletion of *tapA* significantly reduced mScarlet fluorescence compared to wild-type cells, confirming its role in activating expression of *pirG*. Furthermore, expression of *mtrB-P214L* partially restored mScarlet expression in Δ*tapAmctB* cells (Figure 5c), a two-fold difference that was not detectable by RT-qPCR despite elevated MtrA∼P levels (Figure 4d-f). These results confirm that expression of *mtrB-P214L* can partially restore expression of the MtrA regulon in Δ*tapAmctB* cells.

To quantitatively analyze expression dynamics and determine the timing of *pirG* expression, we used time-lapse microscopy to monitor fluorescence from birth to division in 10 individual cells and their 20 daughter cells grown in HdB media and observed cyclic expression of *pirG* (Figure 6a-b, Figure S7a). Since the reporter protein has a long half-life, fluorescence at the first time point reflects prior gene expression. Thus, we subtracted the initial signal intensity of each mother cell from subsequent timepoints within a lineage. To account for the fluorescent protein’s maturation time (∼40 minutes for mScarlet-I [31]), we also constructed strains expressing mScarlet driven from the *ftsZ* promoter sequence, a gene with known mid-cell-cycle expression in *M. tuberculosis* [32]. This calibration allowed us to temporally align expression patterns and confirmed that *pirG* expression peaks near the end of the cell cycle. In contrast, mScarlet expression from a constitutive promoter derived from the ribosomal gene *rpsA* (P*_uv15_*) [21] was not cyclic or patterned across lineages (Figure 6c, Figure S7a). We also tested the *pirG*, *ftsZ*, and *uv15* reporters in wild-type cells grown in osmo-protective HdB-SMM and observed similar results, indicating that increased osmolarity does not alter the cyclic expression patterns in wild-type cells (Figure S7b).

**Figure 6:**
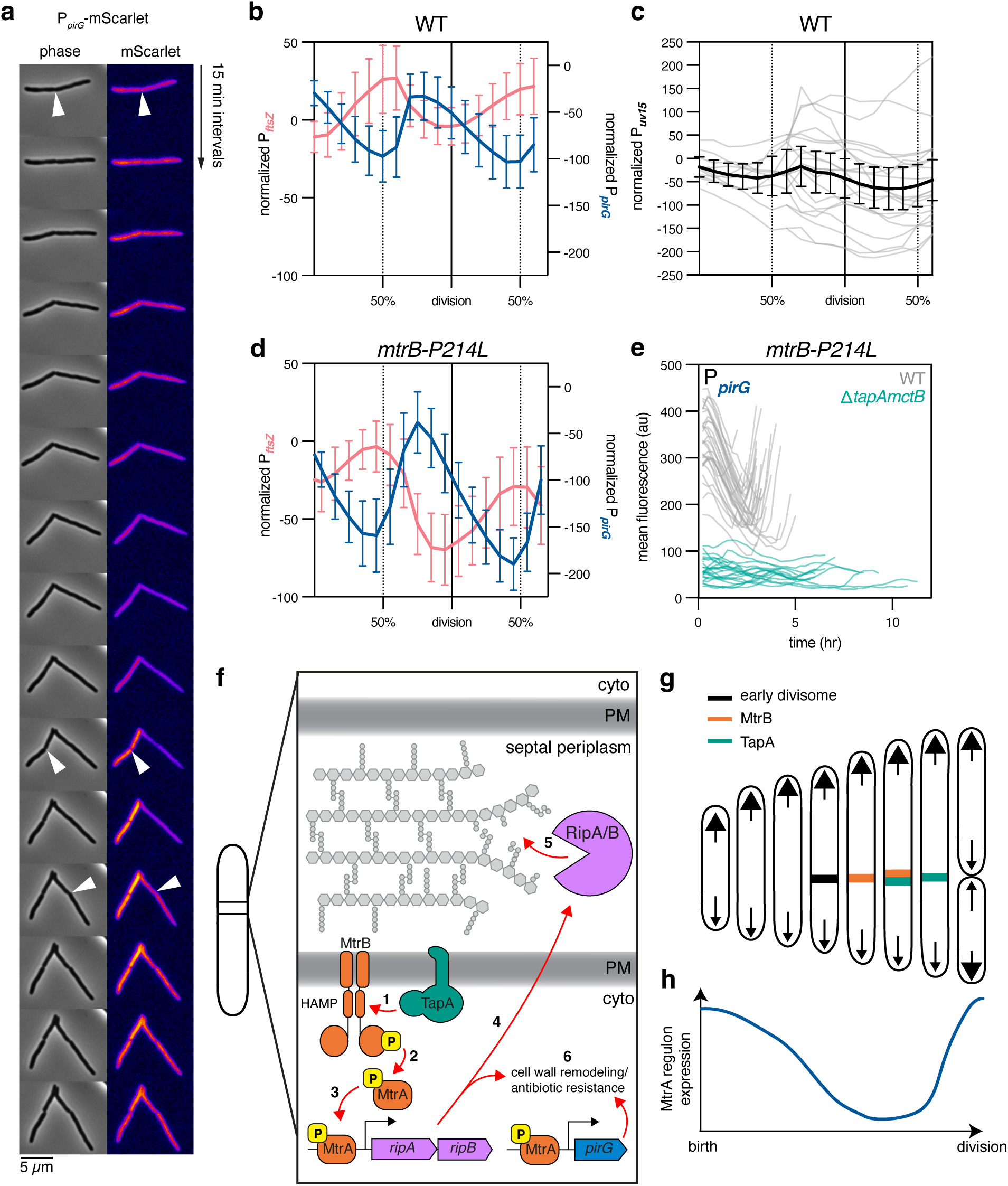
MtrA-mediated gene expression is cell cycle controlled and requires TapA. **a**, Representative wild-type cells expressing mScarlet-I driven by the *pirG* promoter, imaged by time-lapse fluorescence and phase contrast microscopy. Each frame is a 15-minute interval, and white arrow heads indicate cell divisions determined by invagination in the phase contrast channel. **b**, mScarlet-I expression driven by the *ftsZ* promoter or *pirG* promoter in a wild-type strain background across 2 generations. *n* = 10 mother cells; *n* = 20 daughter cells. mScarlet-I signal is normalized to the starting signal of each mother cell within a lineage. Means and 95% confidence interval are shown. **c**, mScarlet-I expression driven by constitutive *uv15* promoter in wild-type cells. Individual cell traces (gray lines) normalized to mother cell are shown. Black line indicates mean and 95% confidence interval. *n* = 10 mother cells; *n* = 20 daughter cells. **d**, In the *mtrB-P214L* background, mScarlet-I expression driven by the *pirG* or *ftsZ* promoter, as in **b**. Means and 95% confidence interval are shown. **e**, mScarlet-I expression from the *pirG* promoter in cells expressing *mtrB-P214L* in either wild-type or *ΔtapAmctB* strain backgrounds. *n* = 30 cells (WT), n = 25 cells (Δ*tapAmctB*). Individual cell traces are shown without normalization. **f-h**, Model of TapA activation of MtrAB and subsequent gene expression. **f**, (1) TapA activates MtrB autophosphorylation. (2) MtrB phosphorylates MtrA. (3) MtrA-P then binds to promoters of target genes such as *ripAB*, and other genes mediating antibiotic resistance, promoting their expression. (4) RipAB proteins are exported to the septal periplasm where they (5) hydrolyze peptidoglycan and separate the daughter cells at the end of division. (6) *ripAB* and *pirG* contribute to cell wall remodeling and subsequent antibiotic resistance **g**, Early divisome proteins, such as FtsZ and associated PG synthases, build the septum near mid-cell. MtrB arrives later in division, followed by TapA. MtrB signal soon dissipates, while TapA remains at the septum until division is complete. **h**, MtrA-mediated gene expression reaches a maximum at the end of division.

Since the mScarlet time-lapse analysis relies on confirmation of division using phase contrast, we were unable to accurately determine the frames at which birth and division occur in chained Δ*tapAmctB* cells. Instead, to directly test the role of TapA in regulating cyclic gene expression, we quantified *pirG* reporter fluorescence in cells expressing *mtrB-P214L* with and without *tapA*. Time-lapse microscopy of *pirG* reporter fluorescence in cells expressing *mtrB-P214L* and *tapA* confirmed that the P214L MtrB mutation increases MtrA regulon expression (Figure 5d, Figure S7c), as evidenced by a greater amplitude of mScarlet expression compared to cells carrying wild-type *mtrB* (Figure 6b). Importantly, deletion of *tapA* abolished the cyclic nature of *pirG* expression (Figure 56e), establishing TapA’s role in driving the dynamic expression of the MtrA regulon. Intriguingly, *ftsZ* expression was also lower and less clearly cyclic (Figure S7d), likely due to indirect dysregulation of cell cycle progression in Δ*tapAmctB* cells, as *ftsZ* is not a known member of the MtrA regulon. Collectively, these data support a model in which TapA mediates the MtrAB-dependent signaling cascade late in the cell cycle, leading to expression of multiple genes, including those involved in cell separation and intrinsic antibiotic resistance (Figure 6f-h).

## DISCUSSION

In *C. glutamicum*, SteA and SteB form a complex at the septum, where SteB directly activates the peptidoglycan hydrolase RipA by removing an autoinhibitory domain from RipA’s active site [26]. Recent studies, however, indicate that SteA may also have functions independent of SteB in *C. glutamicum* [13]. Consistent with these findings, our data demonstrate that in both *M. smegmatis* and *M. tuberculosis*, the putative SteB homolog, MctB, is dispensable for TapA recruitment and subsequent cell separation in typical laboratory conditions. Therefore, TapA’s essential function in mycobacteria is independent of SteB/MctB. Although SteB/MctB is not necessary for cell separation in normal laboratory conditions, it may become essential in specific conditions. For instance, acidic environments activate the protease MarP to cleave RipA’s autoinhibitory domain [33]. Nevertheless, other factors must be responsible for RipA activation under standard conditions. PonA1, a bifunctional peptidoglycan synthase known to bind RipA, may fulfill some aspect of this role [14].

TapA’s essential role in standard culture conditions is to promote the expression of the essential peptidoglycan hydrolases *ripAB* via activation of the two-component system MtrAB. This conclusion is supported by data showing that the essentiality of *tapA* can be bypassed by driving expression of *ripAB* with an MtrA-independent promoter (Figure 2e-f) or with a gain-of-function MtrB allele (Figure 4d-e, Figure 6d-e). However, TapA could have additional non-essential roles *in vitro* or even other essential roles in different environments, such as during infection. Supporting this possibility, our pulldown experiments identified interactions between TapA and other histidine kinases, including MprB and PhoR, the latter of which is strongly activated during infection [28]. Collectively, our results suggest that TapA may be required for numerous gene expression events in *M. tuberculosis*, thereby strengthening its potential as a drug target.

Our data strongly suggest that TapA interacts directly with MtrB. This is based on co-immunoprecipitation and bacterial two-hybrid results, which show an interaction between TapA and MtrB. Additionally, as TapA can interact with a truncated form of MtrB that is missing the kinase domain, the interaction is facilitated by either the transmembrane helices or the HAMP domain of MtrB. Indeed, the predicted Alphafold3 [34] structure of a TapA-MtrB complex predicts that a TapA homodimer could have several contacts with the HAMP domain of MtrB (Figure S3b). Further research will be necessary to fully understand the detailed mechanism by which TapA activates MtrB; one possibility is that TapA stabilizes the active conformation of the MtrB HAMP domain, thereby promoting MtrB autophosphorylation.

Intriguingly, TapA is annotated as a thiamine pyrophosphate kinase domain-containing protein. While the thiamine binding site is missing in a recently determined structure of *C. glutamicum* SteA, the protein does bind GDP and UDP, suggesting possible enzymatic activity [13]. We do not yet know if TapA has enzymatic activity or if that enzymatic activity is required for its function, but it is possible that a cytoplasmic event is required for TapA activation or septal recruitment. Likewise, histidine kinases typically relay information about the extracellular environment to downstream gene expression events, though how or if a periplasmic signal activates MtrB is unknown. One likely periplasmic modulator of MtrB is the lipoprotein LpqB, which is encoded directly downstream of *mtrB* [35]. Together with our data, this suggests the existence of multiple signals, both cytoplasmic and periplasmic, that are required for full activation of the MtrAB signaling cascade, some of which are mediated by TapA.

Compared to dispersed side-wall growth, polar growth presents a distinct set of constraints for bacterial organisms, while also offering opportunities for drug discovery. For example, one key difference is that the timing between septation and elongation must be more tightly controlled in polar-growing organisms, and dysregulation of this step can be lethal. Many genes in the MtrAB regulation are involved in cell cycle progression and cell wall remodeling, including various cell wall synthetic and remodeling enzymes. Intriguingly, only two genes (*ripAB* and *pirG*) in the regulon have depletion/deletion phenotypes that closely resemble depletion of MtrAB/TapA with respect to drug susceptibility (Figure 2a), suggesting that MtrA-regulated expression of these genes is responsible for the increased drug susceptibility associated with depletion of either *mtrAB* or *tapA*. Additionally, at least in *M. tuberculosis*, MtrA controls the expression of DnaA, which is required for the initiation of chromosome replication [20]. Thus, TapA’s arrival at the septum may link the last step of cell division, cell separation, with the first step, DNA replication, in the next generation.

Cell-cycle-dependent gene expression has been extensively characterized in model bacteria, notably *Caulobacter crescentus* [36–38]. In contrast, the dynamics and importance of such regulation in bacterial pathogens, such as *M. tuberculosis,* remain largely unexplored, even though many genes are cell-cycle regulated [32]. Additionally, the mechanisms underlying intrinsic antibiotic resistance in *M. tuberculosis* are not fully understood. Our findings demonstrate that at least some factors involved in antibiotic resistance are dynamically expressed during the mycobacterial cell cycle—a phenomenon obscured by population-level analyses. Interventions capable of disrupting this dynamic gene expression could substantially enhance antibiotic susceptibility across the bacterial population, offering new therapeutic strategies for tuberculosis treatment.

## ACKNOWLEDGMENTS

This work was supported by NIH R01AI148255 (E.H.R), a Harvey L. Karp Postdoctoral Fellowship (S.L.), the Potts Memorial Foundation (S.L.), and an NIH/NIAID New Innovator Award (1DP2AI144850-01, J.M.R.). The funders had no role in study design, data collection and analysis, decision to publish, or preparation of the manuscript.

## METHODS

### Bacterial strains

*M. tuberculosis* strains are derived from H37Rv. *M. smegmatis* strains are derived from mc2-155. *E. coli* BACTH strains are derived from BTH101.

### Mycobacterial cultures

*Mtb* strains were grown at 37°C in in Difco Middlebrook 7H9 supplemented with 0.2% glycerol, 0.05% Tween-80, 1x oleic acid-albumin-dextrose-catalase (OACD) and appropriate antibiotics (kanamycin 10-20 µg mL). Anhydrotetracycline (ATc) was used at 100-200 ng/mL. *Mtb* cultures were grown standing in tissue culture flasks (unless otherwise indicated) with 5% CO_2_.

*M. smegmatis* strains were grown at 37°C in minimal Hartmans-de Bont (HdB) media made without CuSO_4_. HdB contains 27.4mM glycerol, 15.1mM (NH_4_)_2_SO_4_, 3.42µM EDTA, 49.19µM MgCl_2_ hexahydrate, 0.68µM CaCl_2_ dihydrate, 0.08µM Na_2_MoO_4_ dihydrate, 0.17µM CoCl_2_ hexahydrate, 0.62µM MnCl_2_ tetrahydrate, 0.70µM ZnSO_4_ heptahydrate, 1.80µM FeSO_4_ heptahydrate, 0.89mM Na_2_HPO_4_, and 0.71mM KH_2_PO_4_, and 0.05% (w/v) Tween-80. Where specified, HdB was supplemented with sucrose-magnesium chloride-maleate (HdB-SMM). A 10x stock SMM solution was prepared containing 1M sucrose, 40mM MgCl_2_, and 0.04M maleate, and is pH adjusted to 7. HdB-SMM was made by diluting 10x SMM 1/10 in HdB.

### Generation of deletion mutants and complementation strains in *M. smegmatis*

Strains used in this study are listed in Table S6, plasmids are listed in Table S7, and primers are listed in Table S8. Mutant strains were constructed by RecET recombineering [11], and complementation plasmids were constructed by PCR amplifying relevant genes with their native promoter sequences into L5-or Tweety-integrating vectors by isothermal assembly. Tags and point mutants were introduced by subsequent PCR and isothermal assembly. Electrocompetent cell were prepared and electroporation was performed as in ref. 10.

### Generation of individual CRISPRi and CRISPRi-resistant complementation strains in *M. tuberculosis*

Individual CRISPRi plasmids were cloned as described previously using Addgene plasmid 166886 [39]. Briefly, the CRISPRi plasmid backbone was digested with BsmBI-v2 (NEB R0739L) and gel purified. sgRNAs were designed to target the non-template strand of the target gene ORF. For each individual sgRNA, two complementary oligonucleotides with appropriate sticky end overhangs were annealed and ligated (T4 ligase NEB M0202M) into the BsmBI-digested plasmid backbone. Successful cloning was confirmed by Sanger sequencing. Individual CRISPRi plasmids were then electroporated into Mtb. Electrocompetent cells were obtained as described in ref. 10. Briefly, an Mtb culture was expanded to an OD600 = 0.8–1.0 and pelleted (4,000 x g for 10 min). The cell pellet was washed three times in sterile 10% glycerol. The washed bacilli were then resuspended in 10% glycerol in a final volume of 5% of the original culture volume. For each transformation, 100 ng plasmid DNA and 100 μl electrocompetent mycobacteria were mixed and transferred to a 2 mm electroporation cuvette (Bio-Rad 1652082). Where necessary, 100 ng plasmid plRL19 (Addgene plasmid 163634) was also added. Electroporation was performed using the Gene PulserX cell electroporation system (Bio-Rad 1652660) set at 2,500 V, 700 Ω and 25 μF. Bacteria were recovered in 7H9 for 24 h. After the recovery incubation, cells were plated on 7H10 agar supplemented with the appropriate antibiotic to select for transformants. To complement CRISPRi-mediated gene knockdown, synonymous mutations were introduced into the complementing allele at both the protospacer adjacent motif (PAM) and the seed sequence (the 8–10 PAM-proximal bases at the 3’ end of the sgRNA targeting sequence) to prevent sgRNA targeting. Silent mutations were introduced into Gibson assembly oligos to generate ‘CRISPRi-resistant’ alleles. Complementation alleles were expressed from the endogenous or hsp60 promoters in a Tweety or Giles integrating plasmid backbone, as indicated in each figure legend and/or the plasmid descriptions. These alleles were then transformed into the corresponding CRISPRi knockdown strain, with the plRL40 Giles Int expressing plasmid where necessary. The full list of CRISPRi and complementation plasmids can be found in Table S7.

### FDAA Staining and Quantification of Cell Area and Septum Number in *M. smegmatis*

Exponential phase cultures were stained with the fluorescent D-amino acid RADA (Tocris #6649) at a final concentration of 20mM for 1 hour at 37°C with agitation. Cells were then pelleted by centrifugation and resuspended in phosphate buffered saline with 0.05% Tween-80 and 4% paraformaldehyde (PBST-PFA) to fix cells and prevent outgrowth of stain. Cells were incubated in PBST-PFA at room temperature for 15 minutes, pelleted, then resuspended in PBST alone. 2µL aliquots were then spotted on 50mm glass bottom dishes (MatTek #P50G-1.5-30-F) and covered in a 1% agarose PBS pad. Cells were imaged using a 60x 1.4 NA Plan Apochromat phase contrast objective (Nikon). Fluorescence was excited using the Spectra X Light Engine (Lumencor), separated using single-or multi-wavelength dichroic mirrors, filtered through single bandpass emission filters, and detected with an sCMOS camera (ORCA Flash 4.0). RADA dye was captured by using a custom TRITC filter system (Ex: 550/15; Em: 595/44) with an excitation time of 50 ms. Area of RADA-stained cells was quantified in the open-source image analysis software Fiji by manual segmentation, and septum number was counted by eye using the RADA stain.

### FDAA Staining and Quantification of Cell Area and Septum Number in *M. tuberculosis*

*M. tuberculosis* late exponential phase cultures were sub-cultured to OD 0.05 in 7H9 broth containing 200 ng/mL ATc and 25µM NADA-green FDAA (Tocris #6648). 1mL cultures were incubated in 24-well plates at 37°C with constant agitation for 5 days. Cells were then pelleted and resuspended in PBST-PFA and incubated for 2 hours at RT for fixation. Cells were then pelleted and resuspended in PBST alone. Cells were then imaged under PBS agarose pads as described above, using phase contrast and a custom FITC filter system (Ex: 470/24; Em: 515/30) with an excitation time of 200 ms. Cell area was quantified in the open-source image analysis software Fiji by manual segmentation, and number of septa were counted by eye using the NADA-green stain.

### Antibiotic MIC determination in *M. tuberculosis*

MIC assays were performed as previously described [14]. Briefly, all compounds were dissolved in DMSO (VWR V0231) and dispensed using an HP D300e digital dispenser in a 384-well plate format. DMSO did not exceed 1% of the final culture volume and was maintained at the same concentration across all samples. All strain growth was synchronized and pre-depleted in the presence of ATc (100 ng ml−1) for 5 d before assay for MIC analysis. Cultures were then back-diluted to a starting OD580 of 0.05 and 50 µl cell suspension was plated in technical triplicate in wells containing the test compound and fresh ATc (100 ng ml−1). Plates were incubated standing at 37 °C with 5% CO2. OD600 was evaluated using a Tecan Spark plate reader at 10–14 d post-plating and percent growth was calculated relative to the DMSO vehicle control for each strain. IC50 measurements were calculated using a nonlinear fit in GraphPad Prism and are reported in Table S3. For all MIC curves, data represent the mean ± s.e.m. for technical triplicates. Data are representative of at least two independent experiments.

### Antibiotic MIC determination in *M. smegmatis*

Serial 2-fold dilutions of antibiotics in HdB-SMM media were pipetted into a 96-well plate, including a no-drug control*. M. smegmatis* strains were grown overnight in biological triplicate to OD 0.5 and diluted to final OD 0.001, except for a no-cell control for normalization. Plates were incubated at 37°C with shaking for 2 days before OD600 measurements were taken using a Tecan Spark plate reader, normalized to the no-cell control lane, and percent growth was calculated relative to the no-drug control for each strain. IC50 measurements were calculated using a nonlinear fit in GraphPad Prism and are reported in Table S3. Data are representative of at least two independent experiments.

### Suppressor mutant screen in *M. smegmatis ΔtapAmctB* and whole genome sequencing

*M. smegmatis* Δ*tapAmctB* cells were plated for single colonies on LB plates, and 8 individual colonies were inoculated in HdB supplemented with 100mM sucrose. Cultures were propagated in 5mL media, and 30µL of each culture were passaged in fresh media every 1-2 days. After 20 days of sub-culturing, turbid mutants arose. Cultures were allowed to sit for 30 minutes at RT to allow for settling of clumping cells, and turbidly-growing cells from the top of the culture were inoculated into fresh HdB with no supplementation, and passaged daily for 2 additional days. The resulting populations were then plated on LB agar plates for single colonies, and individual colonies were again tested for turbid growth in HdB alone. Individual isolates were RADA stained and imaged by phase contrast and fluorescence microscopy, as described above. One isolate displayed a normal septum number (0-1), while five other isolates remained as chains. Genomic DNA was isolated by phenol-chloroform extraction with RNAse treatment before sending samples to SeqCenter for Illumina sequencing and variant calling.

### Coimmunoprecipitation of msfGFP– and FLAG-tagged proteins in *M. smegmatis*

Coimmunoprecipitation was performed as described previously [9]. *M. smegmatis* cultures were grown to OD 0.6 in 250mL. Cells expressing msfGFP-tagged divisome proteins or PonA1-FLAG were washed with 1x phosphate buffered saline (PBS) once. Pellets were resuspended in 1 ml of 1x PBS with 1.25 mM dithiobis(succinimidyl propionate) and incubated for 30 min at 37°C for cross-linking. After incubation, cells were pelleted at 10,000x *g* for 5 min at room temperature and the supernatant was discarded. The pellet was resuspended in lysis buffer (50 mM Tris-HCl, pH 7.4; 150 mM NaCl; 10 ug/ml DNase I; one tablet Roche cOmplete EDTA free protease inhibitor cocktail; and 0.5% Igepal Nonidet P40 Substitute) and lysed with a BeadBug Microtube Homogenizer at 4000 rpm six times for 30 s each, icing in between. Lysed cells were spun down at 15,000x *g* for 15 min at 4°C, and the supernatant was transferred to a clean Eppendorf tube. Lysates were incubated with GFP-Trap Magnetic Agarose (Chromotek), or anti-FLAG magnetic beads (Sigma) and incubated at 4°C overnight, rotating. After incubation, samples were spun down at 2500x *g* for 1 min at room temperature and flow-through was discarded. Beads were washed three times with non-detergent wash buffer (10 mM Tris-HCl, pH 7.4; 150 mM NaCl, 0.5 mM EDTA). Samples were eluted with 2x Laemmli Buffer (Bio-Rad) prepared with 50 mM (DTT) and boiled at 95°C for 5 min, or eluted with 3xFLAG-peptide (Sigma). All samples not already treated with Laemmli Buffer + DTT were prepared for Western blot by addition of Laemmli Buffer + DTT and boiled at 95°C for 5 min, to reverse all cross-links. Western blot validation was performed as described previously [9] with anti-FLAG mouse antibody conjugated to HRP (Sigma A8592) or anti-GFP rabbit primary antibody (Genetex) and goat anti-rabbit conjugated to HRP, and proteins were detected by ECL substrate (ThermoFisher SuperSignal™ West Femto Maximum Sensitivity Substrate, 34094). After verification by Western blot, samples were sent to the Harvard Taplin Mass Spectroscopy Core for in-gel digestion with Trypsin, and analyzed with a Thermo Scientific Orbitrap mass spectrometer. Spectra were determined in-house using the Sequest algorithm and mapped to the *M. smegmatis* proteome.

### Bacterial two-hybrid assays

BTH101 competent *E. coli* cells were co-transformed with vectors encoding T25– and T18-tagged proteins under the control of isopropyl β-D-1-thiogalactopyranoside (IPTG) inducible promoters. For qualitative evaluation of β-galactosidase activity, strains were grown up in LB broth overnight and 3µL were spotted onto selective LB agar plates containing 40µg/mL X-gal and 0.5mM IPTG. Plates were then incubated at 30°C for 24 or 43 hours before photos were taken. For quantitative measurement of β-galactosidase activity, freshly co-transformed BTH101 cells were plated for single colonies on selective LB agar plates containing 0.5mM IPTG. After 43 hours of incubation at 37°C, colonies were scraped from plates in 1mL of sterile water and normalized to OD600 0.5. 100µL of normalized cells were diluted in 900µL Z-buffer (60mM Na_2_HPO_4_, 40mM NaH_2_PO_4_, 10mM KCl, 1mM MgSO_4_, 50mM β-mercaptoethanol, pH adjusted to 7) in a 2mL tube in technical triplicate. Next, 50µL chloroform and 25µL 0.1% SDS were added to the tube, followed by vortexing and incubation at 30°C for 5 minutes. 200µL ONPG buffer (4mg/mL ortho-nitrophenyl-β-galactoside in Z-buffer) was added, tubes were inverted twice to mix, then incubated again at 30°C for 5 minutes. β-galactosidase reactions were quenched with 500 µL of 1M Na_2_CO_3_. 100µL of sample were added to a 96-well plate and OD420 and OD550 were measured. Miller units were calculated with the following formula: (OD420 – 1.75*OD550) / (volume of cells added * OD600 *reaction time) * 1000. Data are representative of at least two independent experiments, with *n* = 3 technical replicates per strain.

### Time-lapse microscopy

An inverted Nikon Ti-E microscope was used for the time-lapse and snapshot imaging. An environmental chamber (Okolabs) maintained the sample at 37°C. Exponentially growing cells were cultivated in an B04 microfluidics plate from CellAsic, continuously supplied with fresh HdB or HdB-SMM medium where indicated, and imaged every 15 min using a 60x 1.4 NA Plan Apochromat phase contrast objective (Nikon). Fluorescence was excited using the Spectra X Light Engine (Lumencor), separated using single-or multi-wavelength dichroic mirrors, filtered through single bandpass emission filters, and detected with an sCMOS camera (ORCA Flash 4.0). Filters are: msfGFP (Ex: 470/24; Em: 515/30); mScarlet (Ex: 550/15x; Em: 595/44). msfGFP-TapA was excited for 200 ms and mScarlet was excited for 300 ms for MtrB-mScarlet or 200 ms for mScarlet reporter experiments.

### Total RNA extraction and RT-qPCR in *M. smegmatis*

*M. smegmatis* cultures were grown to OD600 ∼0.8 in biological triplicate. 2 OD units were transferred to a 15mL conical tube with 2 volumes of RNAprotect (Qiagen), vortexed, and incubated at room temperature for 5 minutes. Cells were then pelleted and supernatants were discarded. Cell pellets were resuspended in RLT Buffer (Qiagen) with 2-mercaptoethanol and homogenized by bead beating with Lysing Matrix B silica beads (MP Biomedicals). RNA extraction was performed using the Qiagen RNeasy Kit (74104). 1µg of RNA was reverse transcribed using SuperScript™ IV Reverse Transcriptase (Invitrogen), and qPCR was performed using iTaq Universal SYBR Green Supermix (Bio-Rad). Expression of *ripA* and *pirG* were normalized to that of *sigA* using the ΔΔCt method, and results are expressed as L2FC of wild-type. Primers were designed using the Primer3 online tool (https://primer3.ut.ee). Primer specificity was confirmed for each validated qPCR primer pair through melting curve analysis.

### Total RNA extraction and RT-qPCR in *M. tuberculosis*

Total RNA extraction was performed as previously described [14]. Briefly, 2 OD600 units of bacteria were added to an equivalent volume of GTC buffer (5 M guanidinium thiocyanate, 0.5% sodium N-lauroylsarcosine, 25 mM trisodium citrate dihydrate and 0.1 M 2-mercaptoethanol), pelleted by centrifugation, resuspended in 1 ml TRIzol (Thermo Fisher 15596026) and lysed by zirconium bead beating (MP Biomedicals 116911050). Chloroform (0.2 ml) was added to each sample and samples were frozen at −80 °C. After thawing, samples were centrifuged to separate phases and the aqueous phase was purified by Direct-zol RNA miniprep (Zymo Research R2052). Residual genomic DNA was removed by TURBO DNase treatment (Invitrogen Ambion AM2238). After RNA cleanup and concentration (Zymo Research R1017), 3 µg RNA per sample was reverse transcribed into complementary DNA (cDNA) with random hexamers (Thermo Fisher 18-091-050) following the manufacturer’s instructions. RNA was removed by alkaline hydrolysis and cDNA was purified with PCR cleanup columns (Qiagen 28115). Next, knockdown of the targets was quantified by SYBR green dye-based quantitative real-time PCR (Applied Biosystems 4309155) on a Quantstudio System 5 (Thermo Fisher A28140) using gene-specific qPCR primers (5 µM), normalized to *sigA* (*rv2703*) and quantified by the ΔΔCt algorithm. All gene-specific qPCR primers were designed using the PrimerQuest tool from IDT (https://www.idtdna.com/PrimerQuest/Home/Index) and then validated for efficiency and linear range of amplification using standard qPCR approaches. Primer specificity was confirmed for each validated qPCR primer pair through melting curve analysis.

### Phos-tag^TM^ SDS-PAGE and Western blot

Overnight *M. smegmatis* cultures were grown to OD600 ∼0.6 and 9mL culture were pelleted. Supernatants were discarded, and cell pellets were resuspended in 300µL cold lysis buffer (Tris-buffered saline [TBS] with Roche protease and phosphatase inhibitors). Resuspended cell pellets were homogenized by bead beating with Lysing Matrix B silica beads (MP Biomedicals) at 4°C. Lysates were pelleted, and protein concentrations of the lysate supernatant were normalized in fresh lysis buffer by A595 using Bio-Rad Protein Assay Dye Reagent Concentrate (5000006EDU) to the lowest sample value. Normalized lysates were then mixed with cold Laemmli sample buffer (Bio-Rad), loaded immediately onto SuperSep Phos-tag^TM^ gels (Fujifilm 192-18001), and run for 1.5 hours at constant 30mA at 4°C. To chelate metal ions before transfer, gels were washed 4×10 min in wash buffer with EDTA (25mM Tris-HCl, 192mM glycine, 10mM EDTA), then 2×10 min in wash buffer without EDTA (25mM Tris-HCl, 192mM glycine). Wet transfer to a PVDF membrane was run for 90 min at 90V in transfer buffer (5% methanol, 25mM Tris-HCl, 192mM glycine) at 4°C. Membranes were blocked in TBST (TBS, 0.05% Tween-20) with 5% milk in for 1 hour at room temperature, then incubated in 1° anti-Myc monoclonal antibody (ThermoFisher MA1-21316) in TBST/ 5% milk overnight at 4°C. Membranes were then washed in TBST before adding 2° goat anti-mouse antibody (ThermoFisher A28177) in TBST/ 5% milk for 1 hour at room temperature. Membranes were then washed in TBST again before adding ECL substrate (ThermoFisher SuperSignal™ West Femto Maximum Sensitivity Substrate, 34094). Membranes were incubated with ECL substrate for 5 minutes at room temperature before imaging chemiluminescence. Intensity of bands was quantified using Fiji, and the percentage of phosphorylated (upper) band was expressed as fraction of total protein (lower plus upper bands).

