## Supplemental Figures for "An activator of a two-component system controls cell separation and intrinsic drug resistance in *Mycobacterium tuberculosis*"

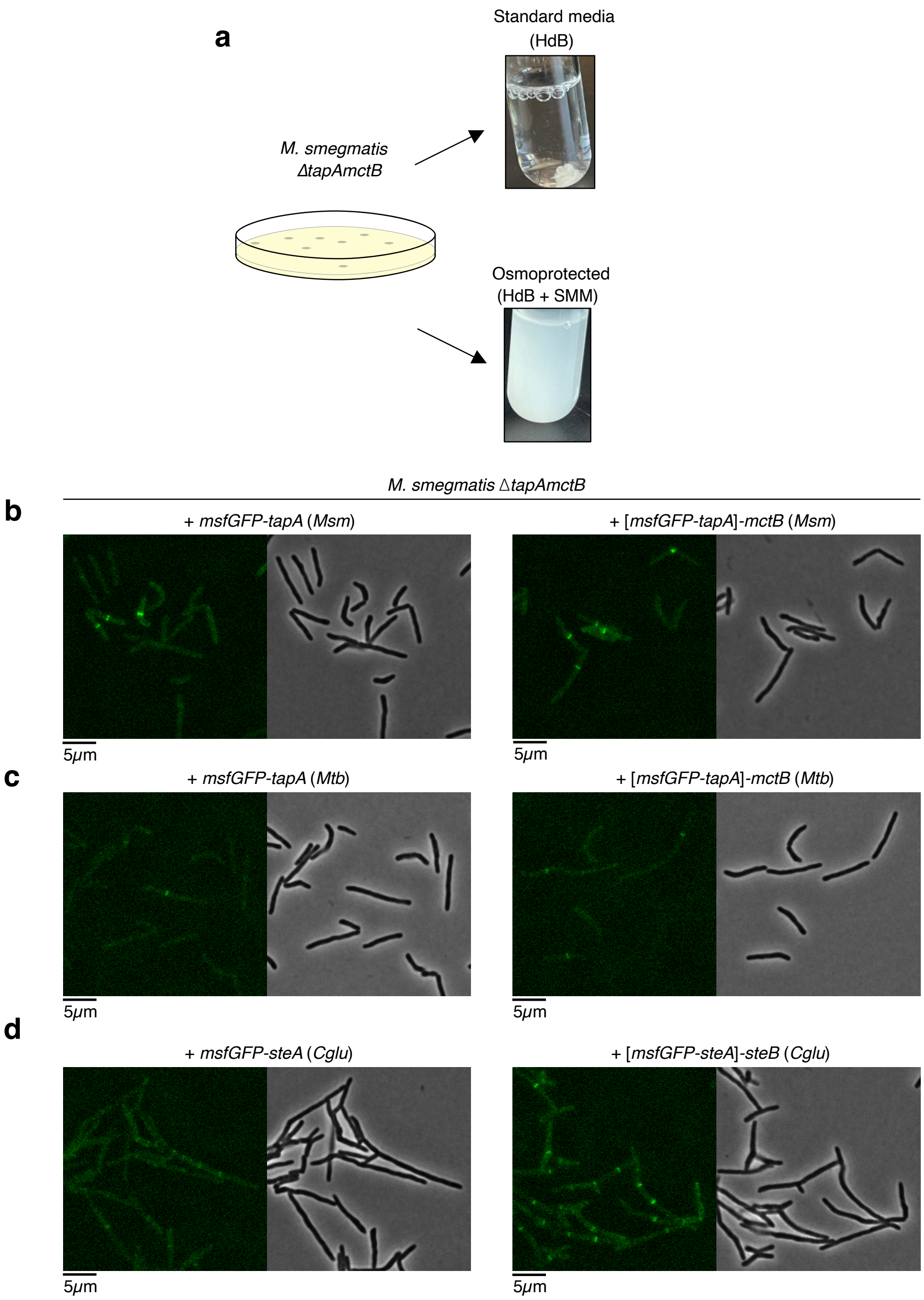


**Figure S1: *M. smegmatis* Δ*tapAmctB* can grow turbidly in SMM-supplemented media, and GFP-tagged *tapA* from *M. smegmatis* or *M. tuberculosis* can complement the mutant. a**, *M. smegmatis* Δ*tapAmctB* transformants grow on selective agar plates, but form clumps that eventually arrest growth when grown in typical HdB media. Supplementation with SMM (sucrose, maleate, magnesium chloride) allows for turbid growth and subsequent passaging. **b-d**, Δ*tapAmctB* complemented with GFP-tagged *tapA*, with or without the corresponding *mctB* homologs from *M. smegmatis* (**b**), *M. tuberculosis* (**c**), or *C. glutamicum* (**d**).

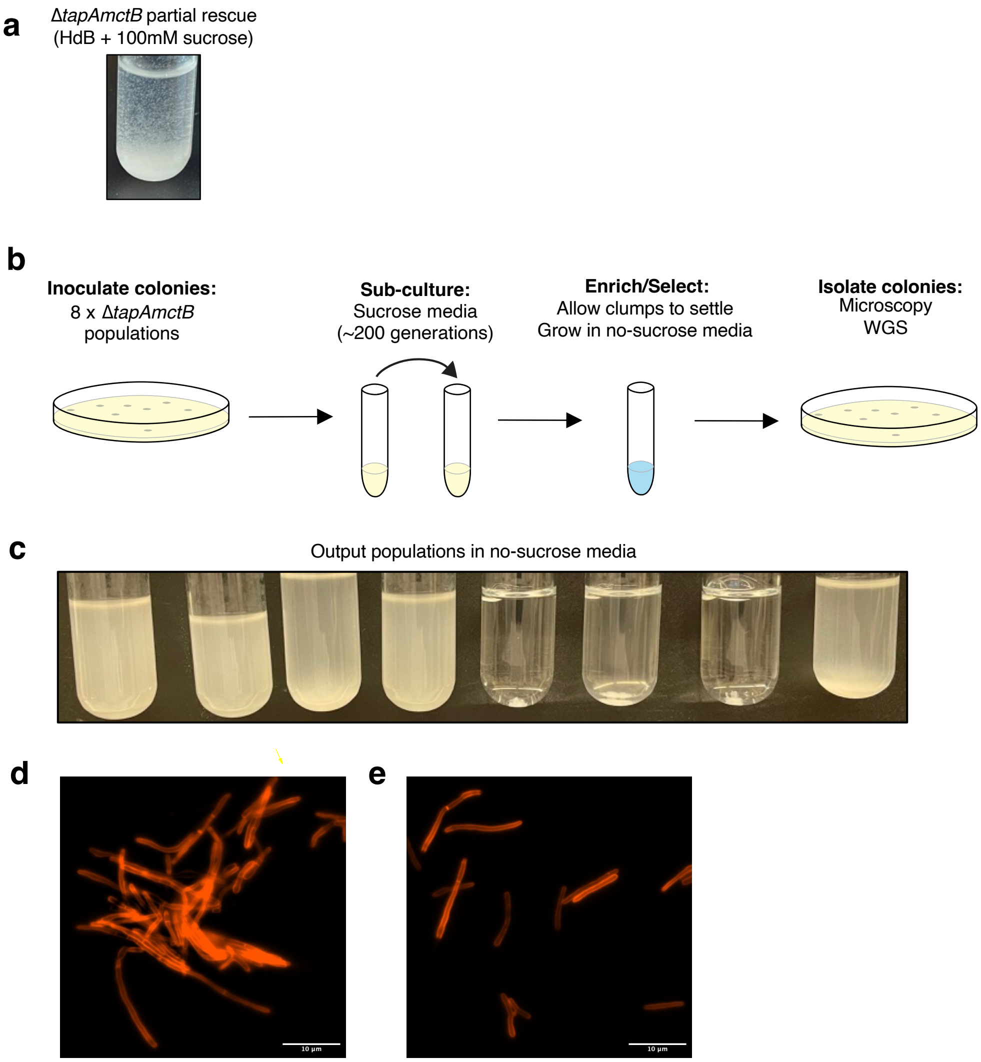


**Figure S2: *M. smegmatis* Δ*tapAmctB* suppressor mutant screen setup. a**, Supplementation of HdB media with 100mM sucrose alone partially rescues Δ*tapAmctB* cells, allowing for propagation and selection of turbidly growing suppressor mutants. **b**, Schematic of screening protocol. Individual Δ*tapAmctB* colonies were inoculated in HdB with 100mM sucrose in 8 parallel culture tubes. 30μL of culture were sub-cultured into fresh HdB-sucrose every 1-2 days for a total of 20 days (~200 generations), after which turbid mutants arose. Turbidly growing cells were then enriched by allowing clumped cells to settle to the bottom of the tube, and then further selected for by sub-culturing in HdB with no sucrose, a non-permissive condition for the original parent strain. **c**, Resulting mixed output populations after growth in no-sucrose HdB. These mixed cultures were then plated on agar to single colonies for phenotypic evaluation and whole genome sequencing of individual isolates. **d-e**, RADA-stained isolates. **d**, One example isolate with no suppression of multiple septa. **e**, The serendipitous mutant suppresses the multiple septa phenotype and contains insertion sequence *IS1549* within the promoter of *ripAB*.

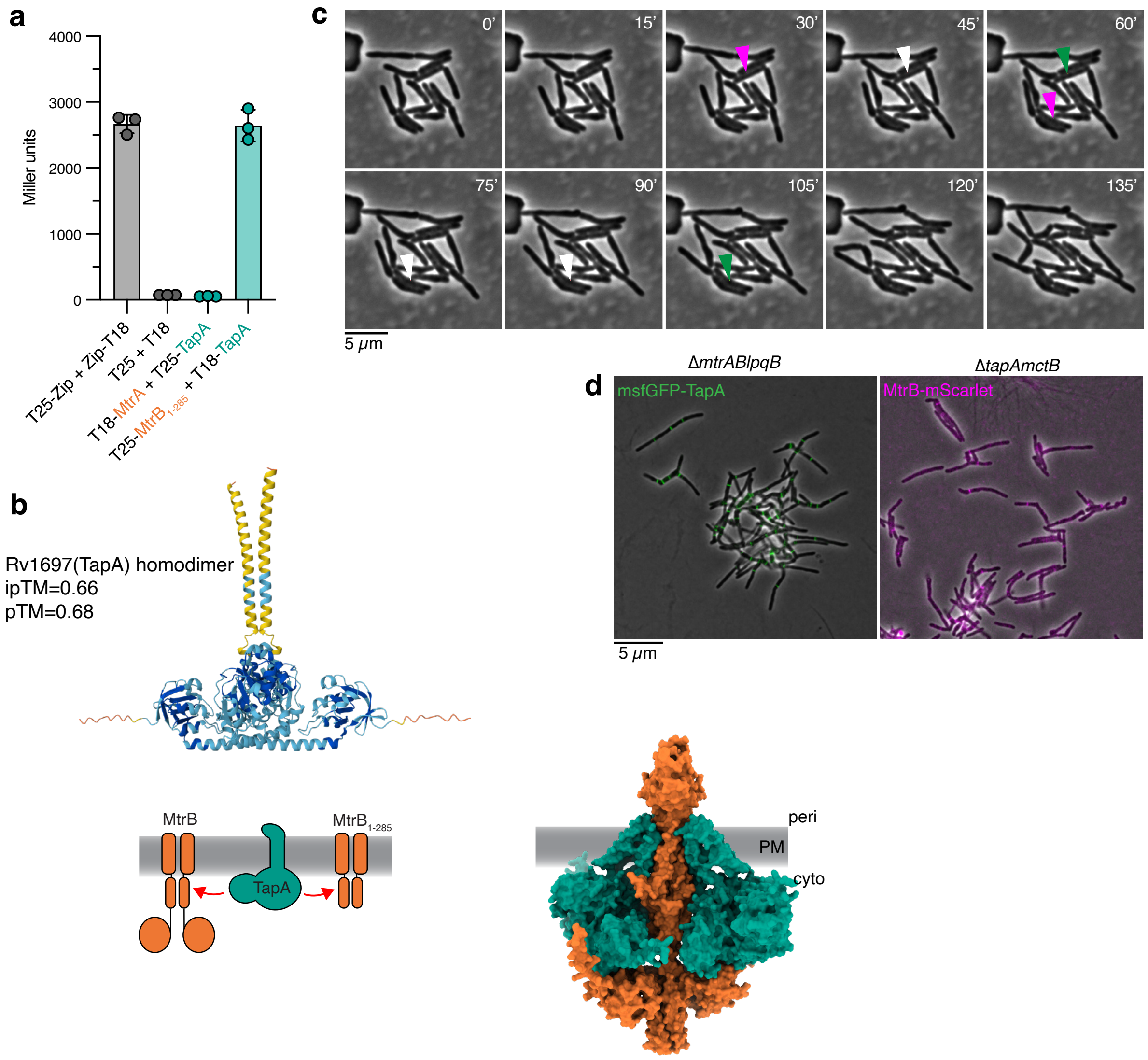


**Figure S3**: **Extended data for Figure 3. a**, BACTH ONPG assay for strains expressing MtrA and TapA or truncation mutant $TapB_{1-285}$ and TapA. **b**, TapA can interact with MtrB or $_{MtrB1-285}$ (top). AlphaFold3 co-folded with two copies of MtrB and four copies of TapA (bottom). **c**, Phase contrast corresponding to fluorescence micrographs in Figure 3B. Colored arrowheads indicate where MtrB-mScarlet (magenta), msfGFP-TapA (green), or both (white) localize in Figure 3. Scalebar = 5μm. **d**, Δ*mtrABlpqB* strain expressing msfGFP-TapA (left) and Δ*tapAmctB* strain expressing MtrB-mScarlet. Overlay of phase contrast and appropriate fluorescence channels. Scalebar = 5μm.

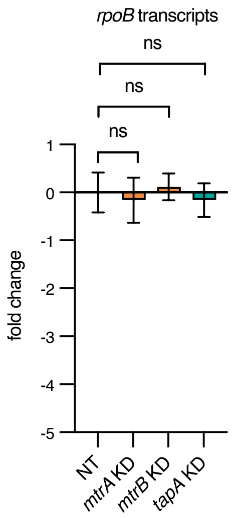


**Figure S4: Expression of *rpoB* is unaltered by knockdown of *mtrA*, *mtrB*, or *tapA* in *M. tuberculosis.*** RT-qPCR as described for Figure 4A.

```
MtrB homologs
MSMEG_1875            199 LLAAIALVVARQIVQPVRSASRIAERFAEGHLTERMPV--RGEDDMARLAVSF-------------- 249
MMAR_1300             225 LLAGIALLVSRQVVVPVRSASRIAERFAEGHLSERMPV--RGEDDMARLALSF-------------- 275
ML0774                225 LLSGIALLVSRQVVVPVRSASRIAERFAEGHLSERMPV--RGEDDMARLAVSF-------------- 275
Rv3245c               225 LLAGIALLVSRQVVVPVRSASRIAERFAEGHLSERMPV--RGEDDMARLAVSF-------------- 275
Mb3273c               225 LLAGIALLVSRQVVVPVRSASRIAERFAEGHLSERMPV--RGEDDMARLAVSF-------------- 275
Clgu0775_ATCC13032    182 LLVGIAWLATQQVTAPVRSASRIAERFAQGKLRERMVV--EGEDEMARLAVSF-------------- 232

Other HAMP domains
HitS                   66 ---------IAVLMRPKREAMIWTIIEPIQKIAKGDFSVKIRNEEKYDGEIGVLVKSINDMTDELNA 122
Rv3645                278 ----------MSIADPLRQLRWALSEVQRGNYNAHMQI--YDASELGLLQAGFNDMVRELSERQR-- 330
NarX                  176 ------------LLQPWRQLLAMASAVSHRDFTQRANI--SGRNEMAMLGTALNNMSAE-------- 228
Aer                   206 ------------IVRPIENVAHQALKVATGERNSVEHL--NRSDELGLTLRAVGQLGLMC------- 255
EnvZ                  180 ------------QNRPLVDLEHAALQVGKGIIPPPLRE--YGASEVRSVTRAFNHMAAGV------- 232
```

**Figure S5: Diverse HAMP domains contain a conserved proline.** Clustal Omega sequence alignment of MtrB homologs and other HAMP-domain containing proteins in other bacteria. MtrB homologs are from *M. smegmatis* (MSMEG_3748), *M. marinum* (MMAR_1300), *M. leprae* (ML0774), *M. tuberculosis* (Rv3245c), *M. bovis* (Mb3273c), and *C. glutamicum* ATCC13032 (Cglu0775). Other HAMP domains are from *Bacillus anthracis* (HitS, a histidine kinase), *M. tuberculosis* (Rv3645, a putative phosphatase), and *E. coli* (NarX, Aer, and EnvZ histidine kinases).

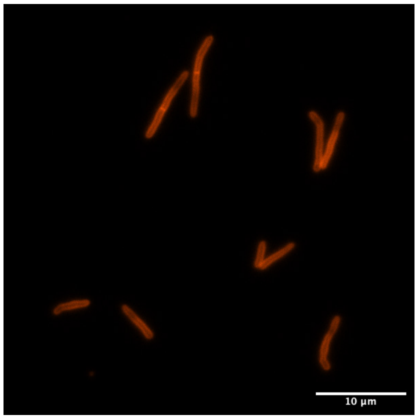


.

**Figure S6: Myc-tagged MtrA complements a Δ*mtrABlqpB* strain.** RADA stain of strain expressing *myc-mtrA* as the sole copy. Scale bar = 10µm.

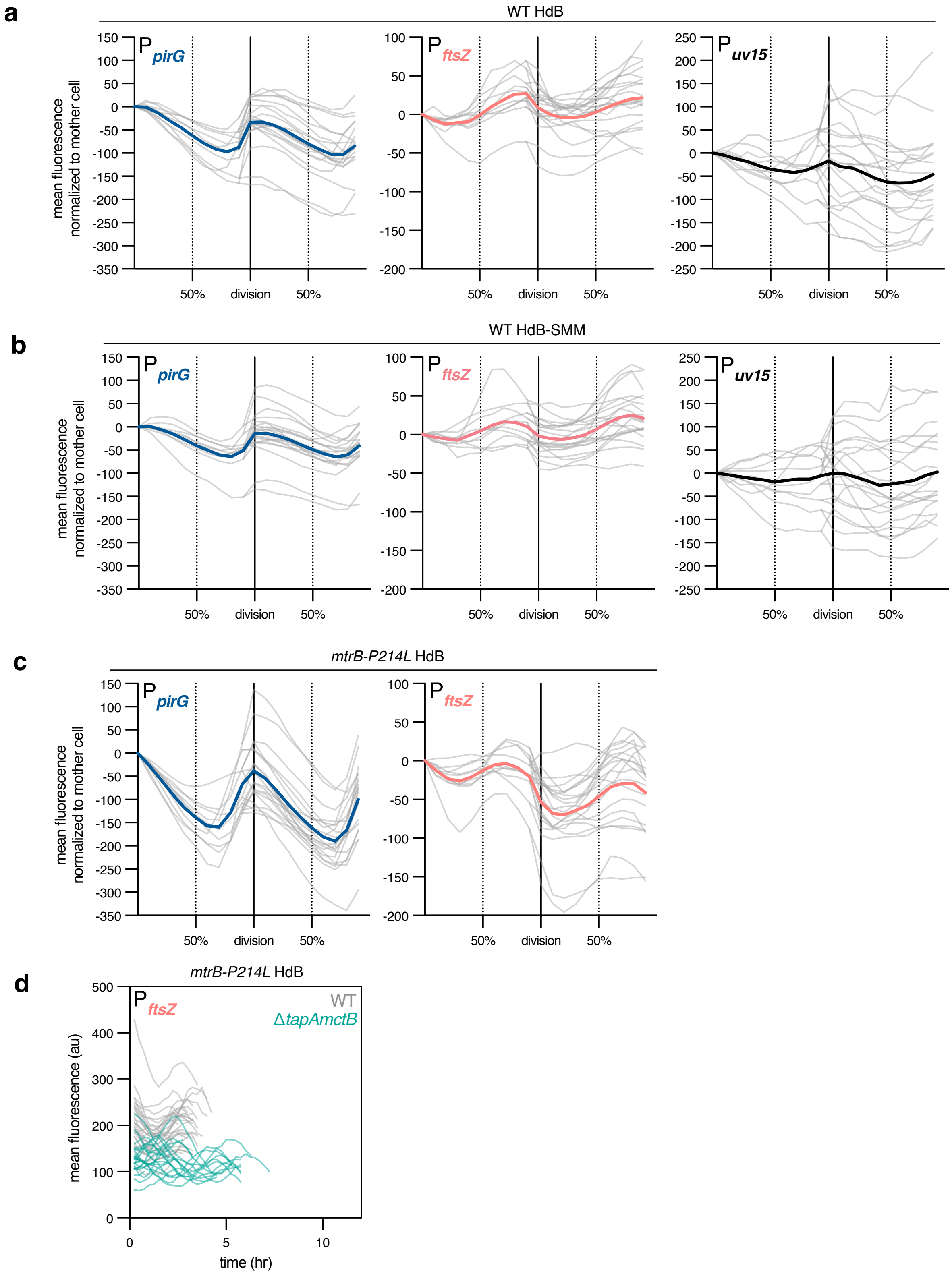


**Figure S7: Individual cell traces and means for mScarlet reporter experiments. a-c**, Single cell traces are shown in gray, and means are showed as thick colored lines as indicated**.** Data are shown without temporal mScarlet correction. **a**, Underlying single-cell traces from Figure 5A-B for wild-type cells grown in HdB with mScarlet driven by the *pirG*, *ftsZ*, or *uv15* promoters. **b**, Single-cell traces for wild-type cells grown in HdB-SMM. **c**, Underlying single-cell traces from Figure 5C for *mtrB-P214L* cells grown in HdB with mScarlet driven by the *pirG* or *ftsZ* promoters. **d**, Single cell traces without cell cycle time normalization for WT or Δ*tapAmctB* strains expressing *mtrB-P214L* and mScarlet driven from the *ftsZ* promoter. Gray lines are WT traces and teal lines are Δ*tapAmctB* traces.
